# Repurposing azithromycin in combination with last-line fosfomycin, colistin and tigecycline against Multi-Drug Resistant *Klebsiella pneumoniae*

**DOI:** 10.1101/2022.07.03.498633

**Authors:** Marta Gómara-Lomero, Ana Isabel López-Calleja, Antonio Rezusta, José A. Aínsa, Santiago Ramón-García

**Affiliations:** Department of Microbiology, Pediatrics, Radiology and Public Health. Faculty of Medicine, and BIFI, University of Zaragoza, Spain; Servicio de Microbiología, Hospital Universitario Miguel Servet, Zaragoza, Spain; CIBER Respiratory Diseases, Carlos III Health Institute, Madrid, Spain; Research & Development Agency of Aragon (ARAID) Foundation, Spain

**Keywords:** antimicrobial resistance, MDR *Klebsiella pneumoniae*, azithromycin, drug repurposing

## Abstract

**Background:** New therapeutical strategies are urgently needed against multidrug-resistant (MDR) Enterobacterales. Azithromycin is a widely prescribed antibiotic with additional immunomodulatory properties, but traditionally underused for the treatment of enterobacterial infections. We previously identified azithromycin as a potent enhancer of colistin, fosfomycin and tigecycline against *Klebsiella pneumoniae* ATCC 13883.

**Objectives:** The aim of this work was to evaluate the antibacterial *in vitro* activity of azithromycin-based combinations with last-line antibiotics against an expanded panel of MDR/XDR *K. pneumoniae* isolates.

**Methods:** Time-kill assays of azithromycin alone and in pair-wise combinations with fosfomycin, colistin and tigecycline were performed against a collection of 12 MDR/XDR *K. pneumoniae* isolates. Synergistic and bactericidal activities of azithromycin-based combinations were analyzed after 8, 24 and 48 hours of treatment, and compared with antimicrobial combinations frequently used in the clinic for the treatment of MDR Enterobacterales.

**Results:** Synergistic interactions were detected in 100% (12/12) for azithromycin/fosfomycin, 58.3% (7/12) for azithromycin/colistin and 75% (9/12) for azithromycin/tigecycline of the strains, showing potent killing activities. Clinical combinations currently used in the clinic showed synergy in 41.6% (5/12) for meropenem/ertapenem, 33.33% (4/12) for meropenem/colistin, 75% (9/12) for fosfomycin/colistin and 66.6% (8/12) for fosfomycin/tigecycline of the strains, with lower bactericidal efficacy.

**Conclusions:** Novel azithromycin-based combinations with last-line MDR/XDR *K. pneumoniae* antibiotics were identified showing *in vitro* capacity to eradicate MDR/XDR *K. pneumoniae*. Our results provide an *in vitro* basis supporting azithromycin used in combinatorial treatment for MDR-related infections.

## INTRODUCTION

Antimicrobial resistance (AMR) is one of the major threats faced by worldwide healthcare systems and, specially, in low- and middle-income countries where the proportion of resistant infections ranges from 40 to 60% compared to 17% for countries belonging to the Organization for Economic Cooperation and Development (OECD)^1^. In 2019, the Center for Disease Control and Prevention (CDC) estimated 210,000 infections and 10,200 deaths in the USA associated to carbapenem-resistant and extended-spectrum beta-lactamases (ESBL)-producing enterobacteria^2^. Among them, carbapenem-resistant *K. pneumoniae* (CRKP) is one of the most concerning superbugs, causing nosocomial infections with mortality rates up to 41.6 and 48%^3^. CRKP incidence is increasing worldwide with 7.9% carbapenem resistance in Europe^4^ and 26.8% of meropenem resistance in China^3^. Moreover, multi-drug resistance is also an increasing trend in *K. pneumoniae*, showing 19.3% combined resistance to traditional first-line antibiotics in the EU^4^.

Although WHO prioritized CRKP as a critical pathogen for antimicrobial development^5^, few new antimicrobial agents are currently in the drug development pipeline; combinatorial therapy with usual antibiotics remains thus the cornerstone therapy for multi-drug resistant (MDR) infections^6,7^. Moreover, the emergence of COVID-19 strongly impacted on AMR; while investment strategies and research advances focused on fighting the virus, disruption of antimicrobial stewardship programs in hospitals have led to an increase of antibiotic misuse^8^, and a rapid spread of resistant bacteria^9^. In this context, drug repurposing (identifying new indications for existing drugs) is an affordable strategy to urgently accelerate the implementation of novel therapies against MDR pathogens^10^.

Azithromycin is a broad-spectrum macrolide antibiotic widely prescribed for several indications such as respiratory, genitourinary and dermal infections^11,12^. Additionally, azithromycin exhibits anti-inflammatory and immunomodulatory properties, demonstrating clinical benefits in critically ill patients^13^ and chronic respiratory disorders such as cystic fibrosis^14,15^, asthma^16^ and chronic obstructive pulmonary disease^17^. This repurposing strategy has been also pursued for azithromycin against parasitic^18,19^ and viral infections^20^. Indeed, azithromycin was one of the first candidates proposed for the management of COVID-19, firstly associated with hydroxychloroquine, although its efficacy for this indication could not be confirmed in clinical trials^21,22^.

Traditionally, monotherapy use of macrolides have been disregarded in the treatment of severe infections caused by Gram-negative bacteria due to different existing mechanisms of resistance to azithromycin in enterobacteria and the low permeability of their outer membrane^23^. However, the enhanced basicity of azithromycin favors the intracellular uptake in Gram-negative bacteria increasing its efficacy and it is currently used for the treatment of enteric infections such as typhoid^12^. In addition, azithromycin’
ss ability to inhibit bacterial quorum-sensing and reducing biofilm formation and mucus production have been demonstrated against intrinsically resistant pathogens (i.e. *Pseudomonas aeruginosa* and *Stenotrophomonas maltophilia*)^24,25^. Moreover, azithromycin therapy seems to exert positive therapeutic effects in murine MDR Gram-negative infection models^26,27^.

In a previous synergy screening, we identified azithromycin as a potent enhancer of last-line antibiotics against MDR enterobacteria^28^. Despite the limitations of azithromycin in monotherapy, its reintroduction into the clinical arsenal to treat high-priority pathogens might be possible in co-administration combination therapy. Here, we evaluated *in vitro* the synergistic and bactericidal activities of azithromycin in combination with fosfomycin, colistin and tigecycline against antibiotic-resistant *K. pneumoniae* isolates and compared them with the activity of combinations typically used in the clinic for the treatment of MDR enterobacteria.

## MATERIALS AND METHODS

### Bacterial strains and growth conditions

A well-characterized set of 12 MDR and Extensively-Drug Resistant (XDR)^29^ *K. pneumoniae* isolates (eight from clinical samples and four from quality assessment exercises) including representative resistance mechanisms was available at the Miguel Servet University Hospital (Zaragoza, Spain) (**Table 1** and **Table S1**). MDR/XDR were defined as: MDR, non-susceptible to ≥1 agent in ≥3 antimicrobial categories; XDR non-susceptible to ≥1 agent in all but ≤2 categories^29^. Bacterial identification was performed by MALDI-TOF mass spectrometry (Bruker Daltonik GmbH, Germany) and antimicrobial susceptibility by an automated broth microdilution method (Microscan Walkaway®, Beckman Coulter, Spain). Phenotypic detection of ESBL, AmpC, carbapenemases and colistin resistance was done according to EUCAST guidelines^30^. Genotypic characterization of resistance mechanisms was performed in clinical samples at the National Microbiology Centre (Majadahonda, Spain). Bacterial LB stocks (15% glycerol) were preserved at -20ºC. Freeze stocks were thawed and sub-cultured on Mueller Hinton broth for 24 hours at 36°C before each assay.

**Table 1.**
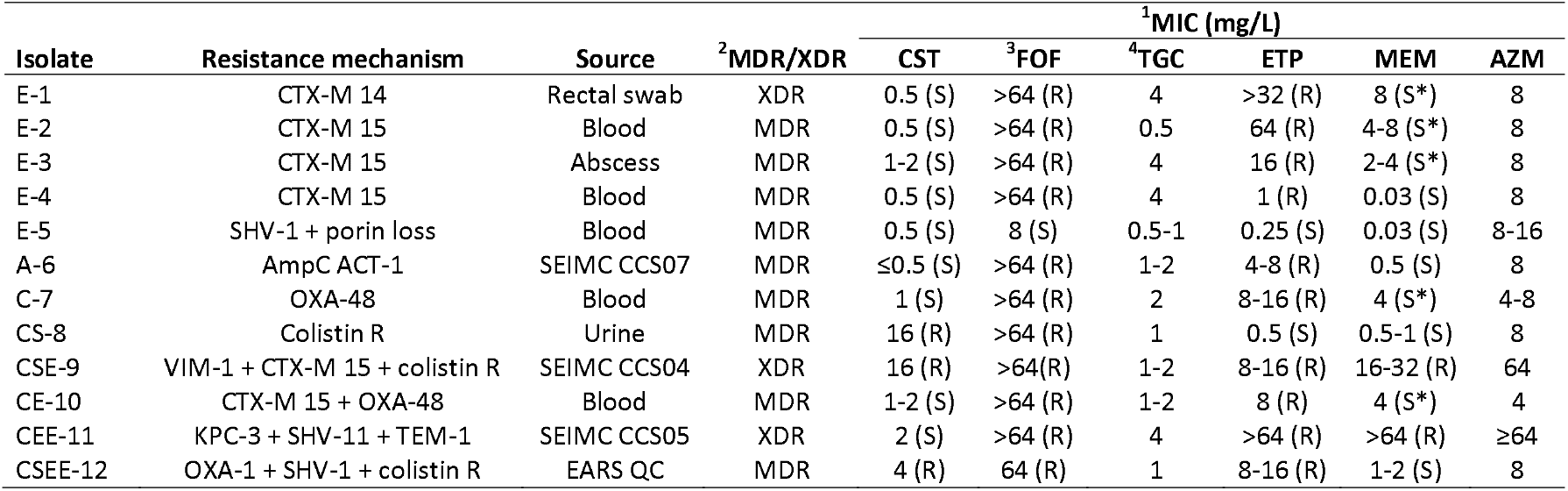
Strain characterization of *K. pneumoniae* isolates and susceptibility profile to drugs used in this study. Clinical categorization according to current EUCAST breakpoints (34) are displayed in brackets. ^1^MIC values were obtained by broth microdilution method in CAMHB. ^2^MDR: non-susceptible to ≥1 agent in ≥3 antimicrobial categories; XDR: non-susceptible to ≥1 agent in all but ≤2 categories (29) (categorization according to susceptibility results provided in Table S1); CST, colistin; FOF, fosfomycin; TGC, tigecycline; ETP, ertapenem; MEM, meropenem; AZM, azithromycin. ^3^The medium was supplemented with 25 mg/L of glucose-6-phosphate for FOF MIC determination ^4^EUCAST clinical breakpoints for tigecycline are only applied to *Escherichia coli* and *Citrobacter koseri* EARS QC, European Antimicrobial Resistance Surveillance Quality Control; R, resistant; S, susceptible; S*: susceptible, increased exposure; SEIMC: Spanish Society of Infectious Diseases and Clinical Microbiology

| Isolate | Resistance mechanism | Source | <sup>2</sup>MDR/XDR | <sup>1</sup>MIC (mg/L) |  |  |  |  |  |
| --- | --- | --- | --- | --- | --- | --- | --- | --- | --- |
|  |  |  |  | CST | <sup>3</sup>FOF | <sup>4</sup>TGC | ETP | MEM | AZM |
| E-1 | CTX-M 14 | Rectal swab | XDR | 0.5 (S) | >64 (R) | 4 | >32 (R) | 8 (S*) | 8 |
| E-2 | CTX-M 15 | Blood | MDR | 0.5 (S) | >64 (R) | 0.5 | 64 (R) | 4-8 (S*) | 8 |
| E-3 | CTX-M 15 | Abscess | MDR | 1-2 (S) | >64 (R) | 4 | 16 (R) | 2-4 (S*) | 8 |
| E-4 | CTX-M 15 | Blood | MDR | 0.5 (S) | >64 (R) | 4 | 1 (R) | 0.03 (S) | 8 |
| E-5 | SHV-1 + porin loss | Blood | MDR | 0.5 (S) | 8 (S) | 0.5-1 | 0.25 (S) | 0.03 (S) | 8-16 |
| A-6 | AmpC ACT-1 | SEIMC CCS07 | MDR | $\leq 0.5$ (S) | >64 (R) | 1-2 | 4-8 (R) | 0.5 (S) | 8 |
| C-7 | OXA-48 | Blood | MDR | 1 (S) | >64 (R) | 2 | 8-16 (R) | 4 (S*) | 4-8 |
| CS-8 | Colistin R | Urine | MDR | 16 (R) | >64 (R) | 1 | 0.5 (S) | 0.5-1 (S) | 8 |
| CSE-9 | VIM-1 + CTX-M 15 + colistin R | SEIMC CCS04 | XDR | 16 (R) | >64 (R) | 1-2 | 8-16 (R) | 16-32 (R) | 64 |
| CE-10 | CTX-M 15 + OXA-48 | Blood | MDR | 1-2 (S) | >64 (R) | 1-2 | 8 (R) | 4 (S*) | 4 |
| CEE-11 | KPC-3 + SHV-11 + TEM-1 | SEIMC CCS05 | XDR | 2 (S) | >64 (R) | 4 | >64 (R) | >64 (R) | $\geq 64$ |
| CSEE-12 | OXA-1 + SHV-1 + colistin R | EARS QC | MDR | 4 (R) | 64 (R) | 1 | 8-16 (R) | 1-2 (S) | 8 |

### Drugs susceptibility testing and media conditions

Azithromycin, fosfomycin disodium salt, glucose-6-phosphate, colistin sulfate, (Sigma–Aldrich, Darmstadt, Germany), tigecycline (European Pharmacopoeia, Strasbourg, France), meropenem (Fresenius Kabi) and ertapenem (MSD) were reconstituted in DMSO or water according to their solubilities. Stock solutions were prepared fresh on the same day of plate inoculation.

Drug susceptibility testing and time-kill assays (TKA) were performed in cation adjusted Mueller Hinton Broth (CAMHB). Minimum Inhibitory Concentration (MIC) determinations were performed by broth microdilution in CAMHB following CLSI guidelines^31^ by the MTT [3-(4,5-dimethylthiazol-2-yl)-2,5-diphenyl tetrazolium bromide] assay^32,33^. Briefly, two-fold serial dilutions of drugs were inoculated with a bacterial suspension of 5×10^5^ CFU/mL in 96-well plates (V_F_= 150 μL) and incubated at 36°C for 18-20 hours. For fosfomycin susceptibility tests, CAMHB was supplemented with 25 mg/L of glucose-6-phosphate, according to EUCAST guidelines^34^. After incubation, 30 μL/well of a solution mix (MTT/Tween 80; 5 mg/mL/20%) were added and plates further incubated for 3 hours at 36ºC. MIC values were defined as the lowest concentration of drug that inhibited 90% of the OD_580_ MTT colour conversion (IC_90_) compared to growth control wells with no drug added.

Minimum Bactericidal Concentration (MBC) was also determined in order to discern bacteriostatic or bactericidal activities. Before MTT addition, 10 μL/well were transferred to 96-well plates containing LB agar and further incubated at 36ºC for 24 hours before addition of 30 μL/well of resazurin; a change from blue to pink indicated bacterial growth. The MBC was defined as the lowest concentration of drug that prevented this colour change. A compound was considered bactericidal when MBC/MIC ≤ 4^32^.

### Time-kill assays

Exponentially growing cultures of *K. pneumoniae* strains were diluted in CAMHB and inoculated in duplicates in 96-well plates (V_F_= 280 μL/well; 5×10^5^ CFU/mL) containing increasing concentrations (0.1x, 0.25x, 1x, 4x, 10x MIC values) of compounds alone, and incubated at 36ºC. Drug-free wells were used as growth controls and MIC of single drugs were performed in parallel with the same inoculum to ensure compound activity. Samples were taken at 0, 2, 5, 8, 24 and 48 hours, and bacterial population was quantified by spot-platting 10-fold serial dilutions onto Mueller Hinton agar (MHA) plates. Plates were incubated overnight at 36°C and CFU/mL calculated. The lower limit of detection was 50 CFU/mL.

The activity of the three-novel azithromycin-based combinations (fosfomycin/azithromycin, colistin/azithromycin and tigecycline/azithromycin) was compared with that of four usual MDR clinical treatments (meropenem/ertapenem, meropenem/colistin, fosfomycin/colistin and fosfomycin/tigecycline).

To assess the activity of the combinations, dose-response curves of compounds alone were first analyzed to select appropriate concentrations for combinatorial testing. Then, selected concentrations were used in TKA, as described above.

A synergistic combination was defined as a ≥2 log_10_ CFU/mL decrease in bacterial count compared to the most active single agent in the combination at any 8, 24 and 48 hours. Antagonism was defined as a ≥2 log_10_ increase in CFU/mL between the combination and the most active single agent. All other degrees of interaction were characterized as indifferent. Bactericidal activity was defined when no bacteria could be recovered in the TKA with a limit of detection of 50 CFU/mL^35^.

## RESULTS

### Activity of azithromycin against MDR/XDR *K. pneumoniae* isolates

There are no CLSI or EUCAST guidelines describing azithromycin clinical breakpoints for enterobacteria, except for *Salmonella* Typhi and *Shigella* spp.^34^; thus, there is no clinical basis to classify *K. pneumoniae* isolates as susceptible or resistant strains. We thus performed MIC determinations of azithromycin against our panel of MDR/XDR *K. pneumoniae* isolates and compared them with the activity of other well-established drugs in the treatment of infections caused by MDR *K. pneumoniae*, for which clinical breakpoints do exist. In our experiments, azithromycin exhibited MIC values ranging from 4 to ≥64 mg/L, which were in the same range of values as those epidemiological cut-offs (ECOFFs) stablished by EUCAST for azithromycin in other enterobacteria; for these, confidence intervals range between 4 to 16 mg/L against *Escherichia coli* and between 4 to 64 mg/L against *S. Typhi*^36^. Thus, the number and nature of antibiotic resistance determinants in any of our twelve isolates appeared not to be related with the susceptibility profiles against azithromycin **(Table 1)**.

### Azithromycin-based combinations are more potent *in vitro* than those combinations currently used in the clinic to treat MDR *K. pneumoniae* infections

We previously identified azithromycin as a potent enhancer of colistin, fosfomycin and tigecycline against *K. pneumoniae* ATCC 13883^28^. All three paired combinations displayed a high synergistic and bactericidal profile against the reference strain (see Figure 3 of Gómara-Lomero *et al)*^28^. In order to further characterize the potential antimicrobial activity of azithromycin-based combinations against MDR *K. pneumoniae*, we extended the TKA validation against a panel of twelve MDR/XDR *K. pneumoniae* isolates with representative mechanisms of resistance **(Figure 1)**. At any time-point (8, 24 and 48 hours), synergy rates among currently used combinations for MDR treatment were observed in 41.6% (5/12) for meropenem/ertapenem, 33.33% (4/12) for meropenem/colistin, 75% (9/12) for fosfomycin/colistin, and 66.6% (8/12) for fosfomycin/tigecycline of the isolates tested (**Figure 1** and **Figure S1**). In stark contrast, a high number of synergistic interactions were obtained with azithromycin-based combinations among all isolates (**Figure 1** and **Figure S2**). Notably, this synergistic bactericidal positive interactions in azithromycin-based combinations were observed even when strains displayed a resistant profile to the drugs alone, as in strain CEE-11 (MIC_AZT_ ≥ 64 mg/L; MIC_FOF_ ≥ 64 mg/L) **(Figure 2)**.

**Figure 1.**
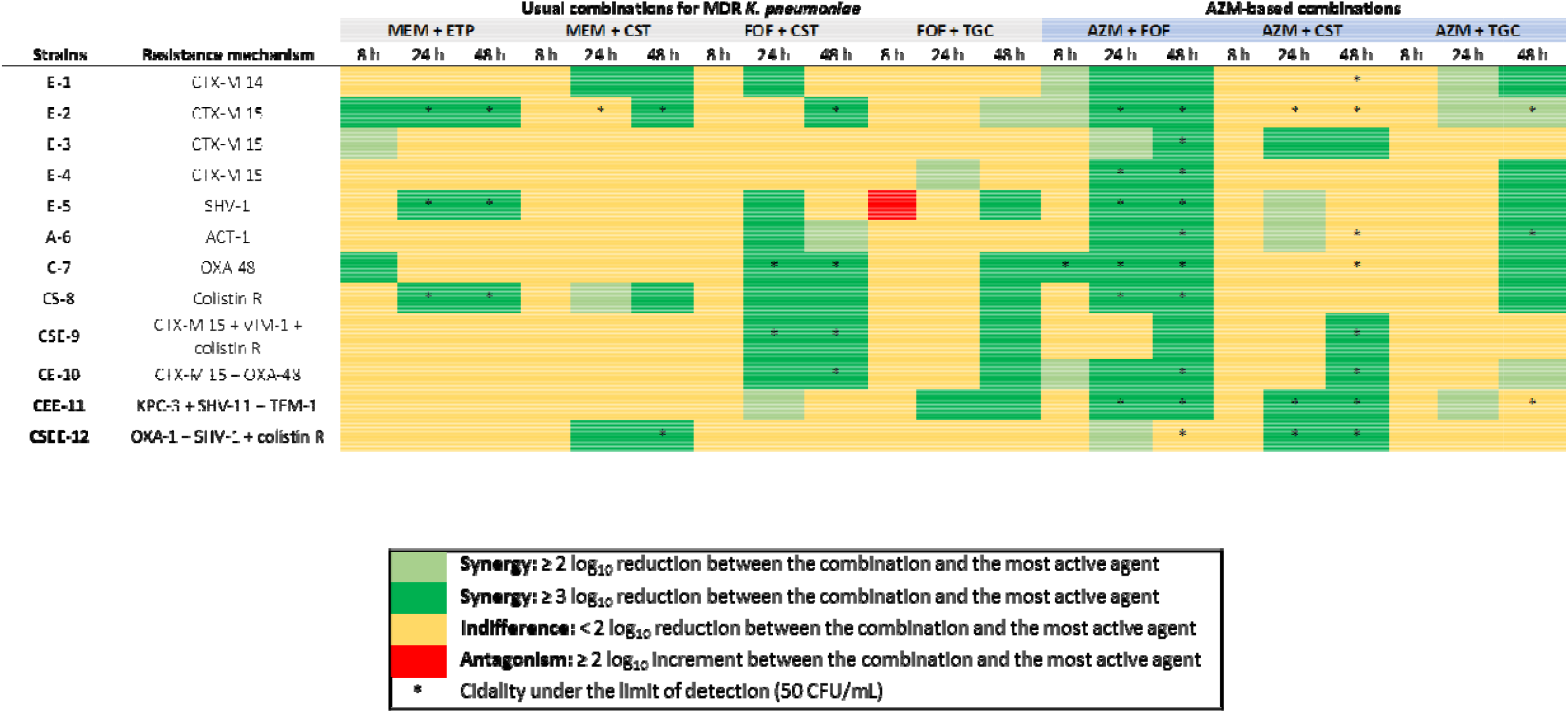
Heat map representation of synergy and bactericidal activities at different time points obtained by time-kill assays against *K. pneumoniae* isolates. Data supporting this summary figure are displayed in Figure S1 and Figure S2. AZM, azithromycin; CST, colistin; ETP, ertapenem; FOF, fosfomycin; MEM, meropenem; TGC, tigecycline.

**Figure 2.**
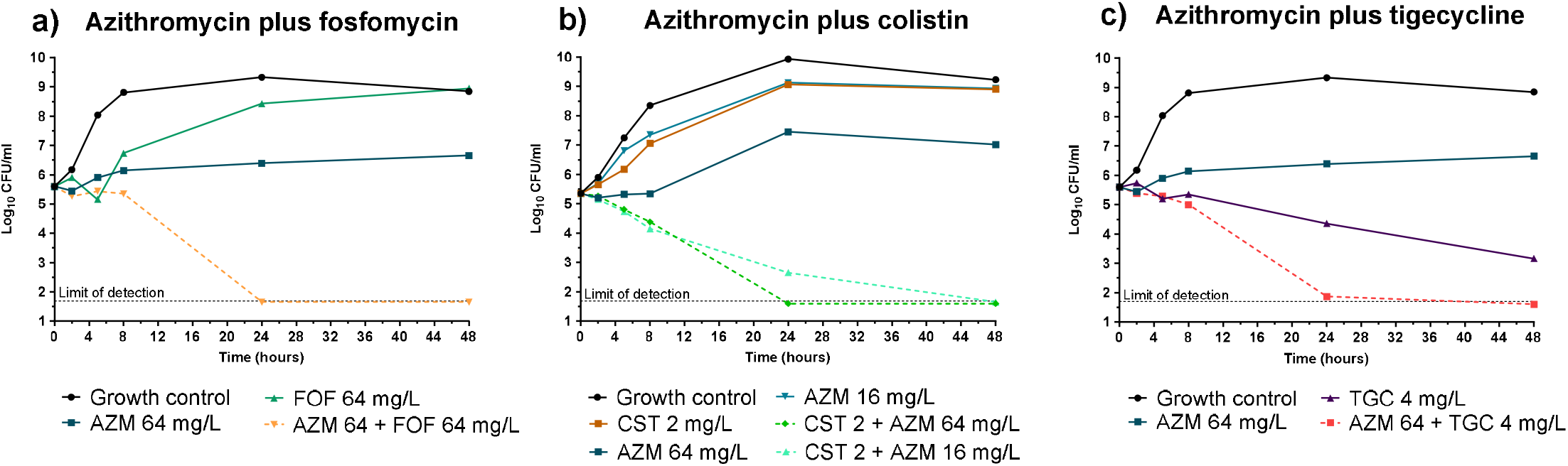
Time–kill curves showing azithromycin combinations with existing antibiotics (a-c) against the *K. pneumoniae* XDR strain CEE-11 (*bla*_KPC-3_ + *bla*_SHV-1_ + *bla*_TEM-1_) in CAMHB. Azithromycin enhanced the activities of fosfomycin, colistin and tigecycline even at subinhibitory concentration (0.25 to 1 x MIC), showing potent synergistic and bactericidal effects. MIC_AZM_ ≥ 64 mg/L, MIC_CST_= 2 mg/L, MIC_FOF_> 64 mg/L, MIC_TGC_= 4 mg/L.

The combination azithromycin/colistin **(Figure S2b)** was synergistic in 7 out of 12 strains (58.3%) and bactericidal in 10 out of 12 strains (83.3%). The positive interaction of azithromycin in combination with colistin was evident when analysing the bactericidal activity at the 48-hour time point in which eight strains (E-1, E-2, A-6, C-7, CSE-9, CE-10, CEE-11, CSEE-12), including two colistin-resistant strains, had viable counts below the limit of detection (50 CFU/mL) **(Figure 1)**.

The combination azithromycin/tigecycline **(Figure S2c)** showed synergistic interactions against 9 out the 12 (75%) strains with a strain-dependent activity. The combination was bactericidal to the limit of detection in three strains (E-2, A-6 and CEE-11) and showed a bacteriostatic profile in the rest of the strains (from <1 to 1.6 log_10_ decrease in CFU/ml), except for CE-10 and CSEE-12 (> 2 log_10_ decrease in CFU/mL at 48 hours) **(Figure S2c)**.

The combination of azithromycin plus fosfomycin was the most potent. This combination was synergistic against all isolates and bactericidal in 11 out of the 12 (91.66%) strains. The potency of the azithromycin/fosfomycin combination was evident when compared to the activity of the drugs alone; neither showed long-lasting bactericidal activity, with a static effect or no activity (azithromycin), and rapid bactericidal activity followed by bacterial regrowth from the 8-hour time-point (fosfomycin). In addition, in most strains combined bactericidal effects were already detected at early time points (4-8 hours) **(Figure S2a)**.

## DISCUSSION

In the present study we evaluated the *in vitro* efficacy of azithromycin in combination with colistin, fosfomycin and tigecycline (currently used last-line antibiotics in the treatment of infections caused by MDR enterobacteria) against a panel of 12 MDR/XDR *K. pneumoniae* isolates with representative resistance patterns. We used TKA as a reference method with activity readouts obtained after up to 48 hours of incubation, a procedure not typically performed when evaluating the activity of compounds against enterobacteria.

We characterized the activity alone of azithromycin, and its three synergistic partners colistin, fosfomycin and tigecycline, in a dose-response manner against our collection of twelve *K. pneumoniae* isolates. Then, we tested them in combination assays selecting matching subinhibitory concentrations of each individual drug to allow for a wider dynamic range and detection of drug interactions. This implies that even if absolute MIC values for every *K. pneumoniae* strain in our collection might be different **(Table 1)**, the effect of their subinhibitory activities would be similar in combination, since they are based on individual MIC values for each strain and compound. The use of subinhibitory concentrations of the antibiotics alone is a key factor to detect drug interactions since higher effective concentrations might masked the effect of their potential interactions. In addition, extending the readout to 48 hours provides information in both the increased bactericidal activity of the azithromycin-based combinations compared to the drugs alone, and also the ability of the combination to completely eradicate bacteria (below the limit of detection of the assay, which is a proxy for culture sterilization). Based on these criteria, we tested three azithromycin-based combinations (**Figure 1** and **Figure S2**) and compared them with four representative combinations currently used in the clinic to treat MDR/XDR *K. pneumoniae* infections (**Figure 1** and **Figure S1**). Our TKA data showed high rates of favourable interactions for the azithromycin-containing combinations, even against strains with concurrent resistance mechanisms; thus, suggesting a potential role of azithromycin in combinatorial therapy (**Figure 1** and **Figure S2**), as evidence by the examples below:

### (i) Azithromycin plus fosfomycin

First prescribed for urinary tract infections, fosfomycin was identified as synergistic partner of several antibiotics. Fosfomycin is an old bactericidal antibiotic that inhibits peptidoglycan synthesis^37^, thus it could be enhancing antibiotic entrance by increasing cell permeability. As such, fosfomycin has been reintroduced in combinatorial therapy for the clinical management of MDR enterobacterial infections over the last years^38^. This combination was previously assessed in two other *in vitro* studies. Presterl *et al*. described negligible bactericidal activity against biofilm-producer *Staphylococcus epidermidis*^39^, and the combination also showed killing activity at 24 hours by TKA against *Neisseria gonorrhoeae*, including azithromycin resistant strains, with no regrowth until the end of the assay^40^. The latter study is in agreement with our results in *K. pneumoniae*, supporting the potential use of azithromycin/fosfomycin against Gram-negative bacteria. We observed rapid bactericidal activities maintained up to the end of the assays against all tested strains (**Figure 1** and **Figure S2a**), including those strains with high fosfomycin MIC values **(Figure 2)**. Interestingly, effective fosfomycin concentrations in our *in vitro* assays were below fosfomycin peak plasma concentration after intravenous administration in adults (606 mg/L)^37^. To the best of our knowledge, this is the first study analyzing the antimicrobial activities of the combination azithromycin/fosfomycin against a large set of MDR *K. pneumoniae* strains. Our results, together with other evidence, suggest that the combination of azithromycin plus fosfomycin could play an important role in clinical settings and merits further pre-clinical and clinical development. Both drugs display good safety profiles, they are recommended for combinatorial therapy to minimize resistance emergence derived from monotherapy, and are administered at a single dose administration (0.5 to 2 g single dose oral or intravenously for azithromycin^12^ and 3 g single dose orally or up to 8 g /8 hours intravenously for fosfomycin^41^). Similar to azithromycin, fosfomycin displays immunomodulatory mechanisms^37^, which have been shown beneficial to overcome severe Gram-negative infections.

### (ii) Azithromycin plus colistin

This combination was reported in some studies including MDR *K. pneumoniae*^26,27,42^, where the increase in the Gram-negative outer membrane permeability facilitates azithromycin access to the 50S ribosomal subunit^26,27^. In agreement with our results, we obtained sterilizing activities in 2 out of 3 of the colistin resistant strains (CSE-9, MIC_CST_= 16 mg/L and CSEE-12, MIC_CST_= 4 mg/L). In these strains, the limiting factor for activity was the concentration of azithromycin; similar killing profiles were obtained at two colistin concentrations (2 mg/L and 8 mg/L) **(Figure S2b)**. These findings support the possibility to decrease colistin concentrations below its nephrotoxic threshold (2.42 mg/L)^43^, if administered in synergistic combination with azithromycin.

### (iii) Azithromycin plus tigecycline

This is the first report of this combination being active against *K. pneumoniae*. Previous studies described biofilm eradication against *S. maltophilia*^25^ and the *in vitro* and *in vivo* activity of azithromycin in combination with minocycline (another tetracycline antibiotic) against MDR pathogens including *K. pneumoniae*^44^. Although we observed variable activity from one strain to another **(Figure S2c)**, the combination showed sterilizing activity against three strains, which had different susceptibility profile to both drugs (e.g., CEE-11 exhibited resistant profile with MIC_TGC_= 4 mg/L and MIC_AZM_ ≥ 64 mg/L, **Figure 2**). Azithromycin and tigecycline are both bacteriostatic drugs targeting the 50S and 30S ribosomal subunits, respectively, which could explain their synergy by enhancing protein inhibition that leads to disruption of the bacterial gene translation.

Azithromycin safety profile is well described, showing uncommon side-effects associated to long-term therapy^45^, and well tolerated when administered to children and pregnant women^46^. It poses advantageous pharmacokinetic and pharmacodynamic (PK/PD) properties respect to other macrolides: no interaction with CYP3A4 cytochrome, an increased tissue penetration and bioavailability due to a higher basic character, and a long half-life (50-70 hours)^11,12^. Peak plasma concentrations of 1.46 mg/L and up to 3.4 mg/L are attained after 1,500 mg-oral and 500 mg-intravenous administrations, respectively^11^. In our study, we observed effective sterilizing activities of azithromycin-based combinations at azithromycin concentrations ranging from 2 up to 64 mg/L **(Figure S2)**. Although for some strains the azithromycin sterilizing concentrations observed were over those achievable in plasma, azithromycin displays a rapid blood-tissue distribution, so despite such low serum concentrations it is expected that its accumulation in tissue will be higher (e.g. accumulation in macrophages is 5-to 200-fold higher than in plasma^12^). In addition, the long post-antibiotic effect and significant subinhibitory concentration effect demonstrated both *in vitro* and *in vivo* against respiratory pathogens^47,48^ indicate a prolonged antimicrobial activity.

The azithromycin PK/PD properties make it an optimal candidate for combination therapy in MDR Gram-negative infections. Standard dosing of the last-line antibiotics used in this study (that included loading doses for colistin and tigecycline)^7^ yielded a rapid bacterial killing effect that could be seconded by the slower but longer lasting action of azithromycin, maintaining bacterial eradication during the course of treatment. Moreover, combinatorial therapy with azithromycin might minimize resistance emergence and toxicity issues (specially with colistin) using longer dosing intervals.

The use of macrolides (specially azithromycin) is currently recommended in critically ill patients with pneumonia as empirical treatment in combination with β-lactams or fluroquinolones^49^, supported by previous preclinical assays showing synergy^50–52^. Anticipatory immunotherapy with azithromycin has been also used in critically ill patients with infections other than pneumonia, demonstrating clinical benefit with reduced mortality rates and intensive-care unit (ICU) stay^13^. The early addition of azithromycin to last-line antibiotics for MDR treatment in severe infections (i.e., sepsis, ventilator-associated pneumonia, immunocompromised patients) could not only improve the efficacy of the therapy in combination, but also improve the clinical outcome due to immunomodulatory properties of azithromycin in ICU patients.

In conclusion, we have demonstrated using *in vitro* TKA models that azithromycin combined with existing antibiotics might increase the efficacy in the eradication of MDR/XDR *K. pneumoniae*. Based on our *in vitro* studies, we propose the following priority list of pairwise combinations: azithromycin/fosfomycin > azithromycin/colistin > fosfomycin/colistin > meropenem/ertapenem > azithromycin/tigecycline > meropenem/colistin > fosfomycin/tigecycline. Additional pre-clinical and clinical studies would be needed to fully understand the clinical potential of azithromycin as synergistic partner in antimicrobial therapies against MDR enterobacteria

## Data availability statement

All data pertaining to this work is within the main manuscript or supplementary information.

## Author Contribution statement

CRediT (Contributor Roles Taxonomy) has been applied for author contribution. Conceptualization, M.G-L. and S.R-G.; Methodology, M.G-L., S.R-G. and A.I.L-C.; Formal analysis, M.G-L.; Investigation, M.G-L. and A.I.L-C.; Resources, A.I.L-C. and A.R.; Data Curation, M.G-L.; Writing - Original Draft, M.G-L., J.A.A. and S.R-G.; Writing - Review & Editing, M.G-L., J.A.A., S.R-G., A.I.L-C. and A.R.; Visualization, M.G-L. and S.R-G.; Supervision, J.A.A. and S.R-G.; Project Administration, S.R-G.; Funding Acquisition, J.A.A. and S.R-G.

## Transparency declarations

None to declare.

