## Supplementary Figures for "Repurposing azithromycin in combination with last-line fosfomycin, colistin and tigecycline against Multi-Drug Resistant *Klebsiella pneumoniae*"

**Figure S1. Time-kill curves against twelve *K. pneumoniae* isolates of pair-wise combinations currently used in the therapy of MDR enterobacteria. (a)** Meropenem plus ertapenem; **(b)** Meropenem plus colistin; **(c)** Fosfomycin plus colistin; **(d)** Fosfomycin plus tigecycline

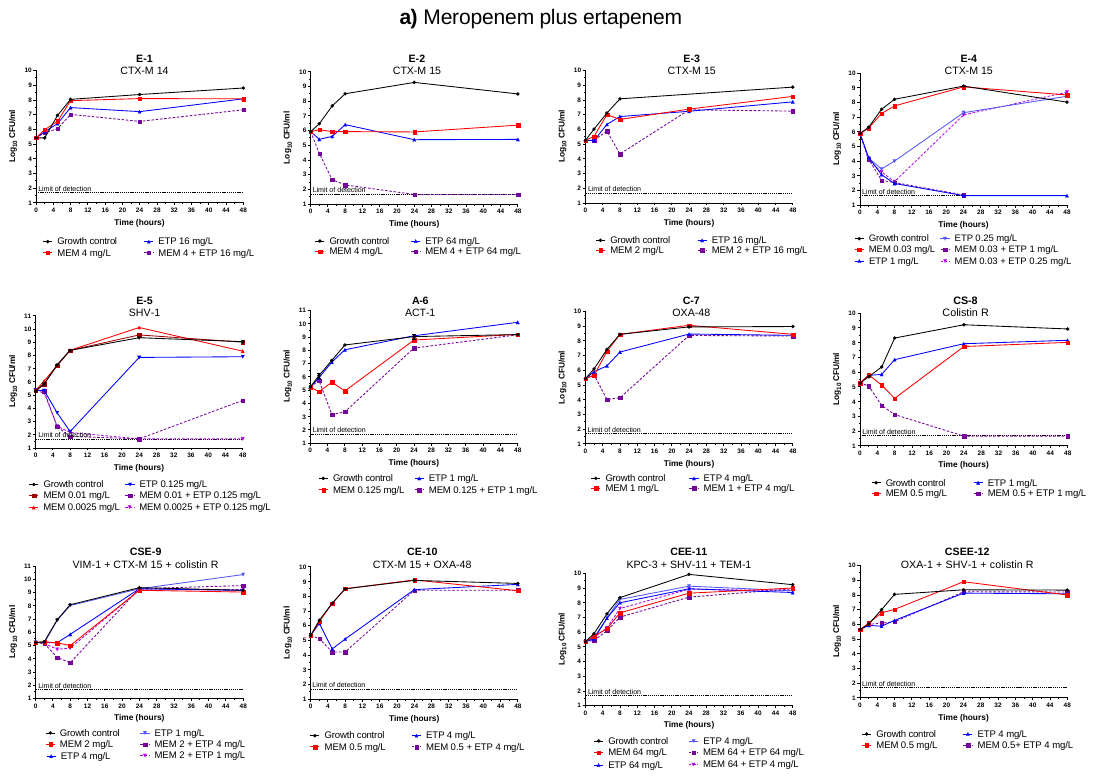

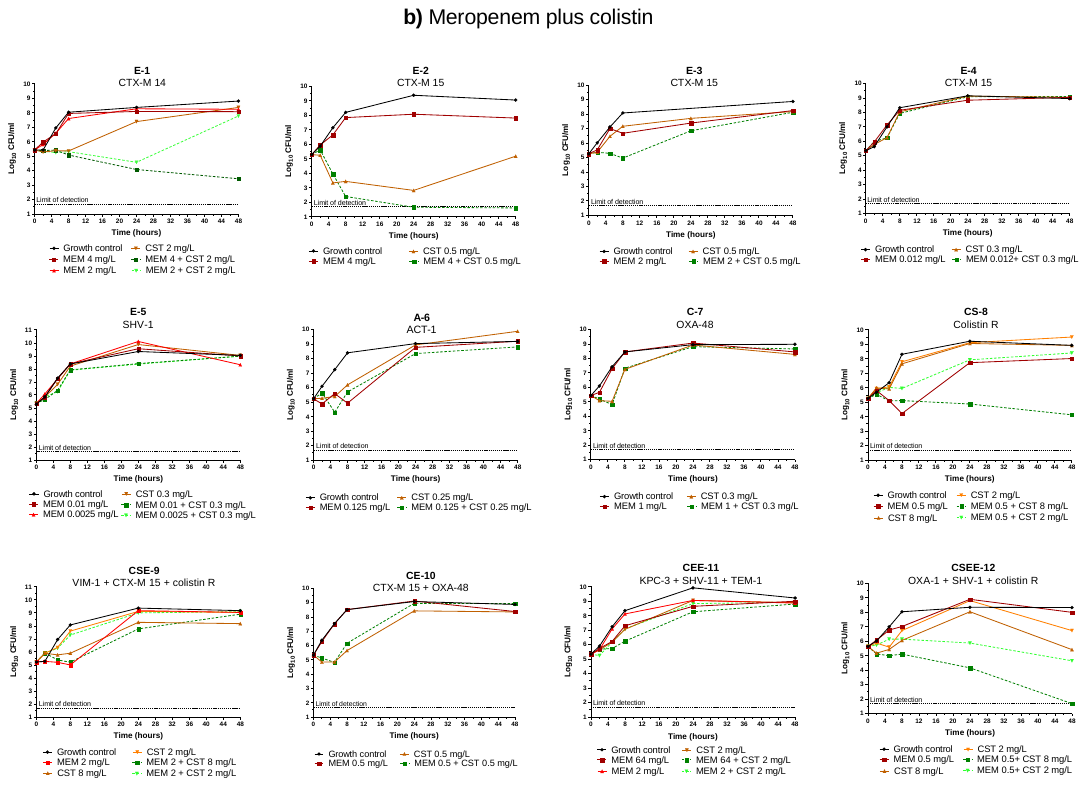

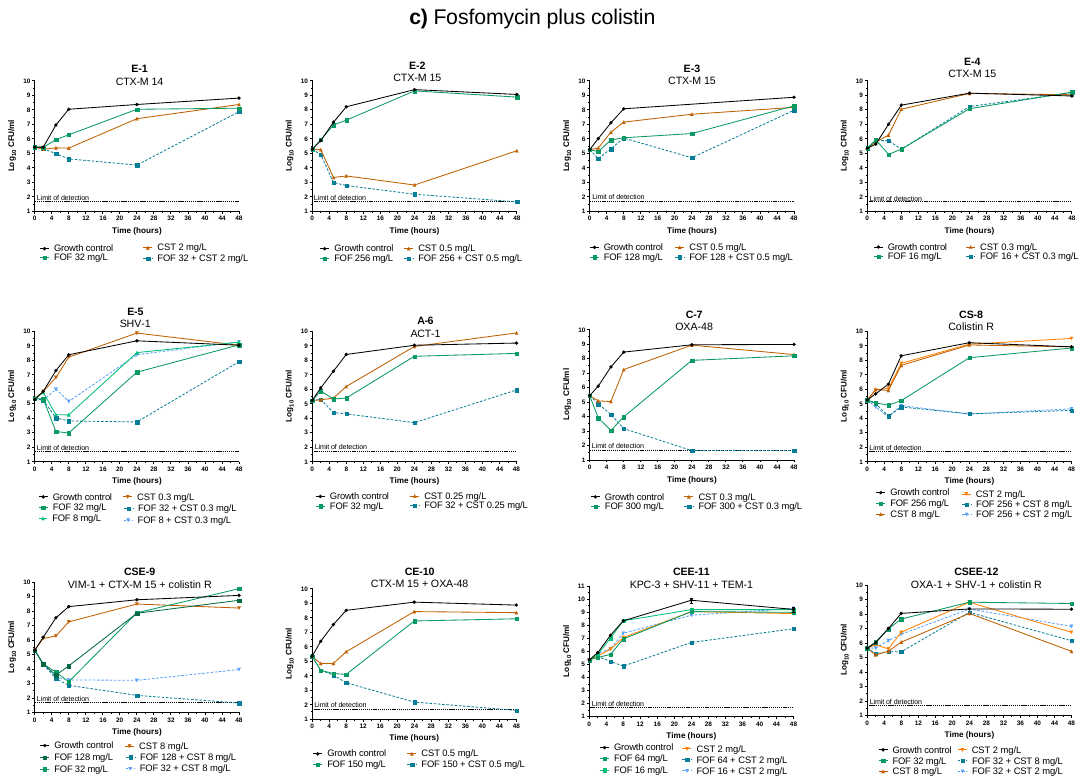

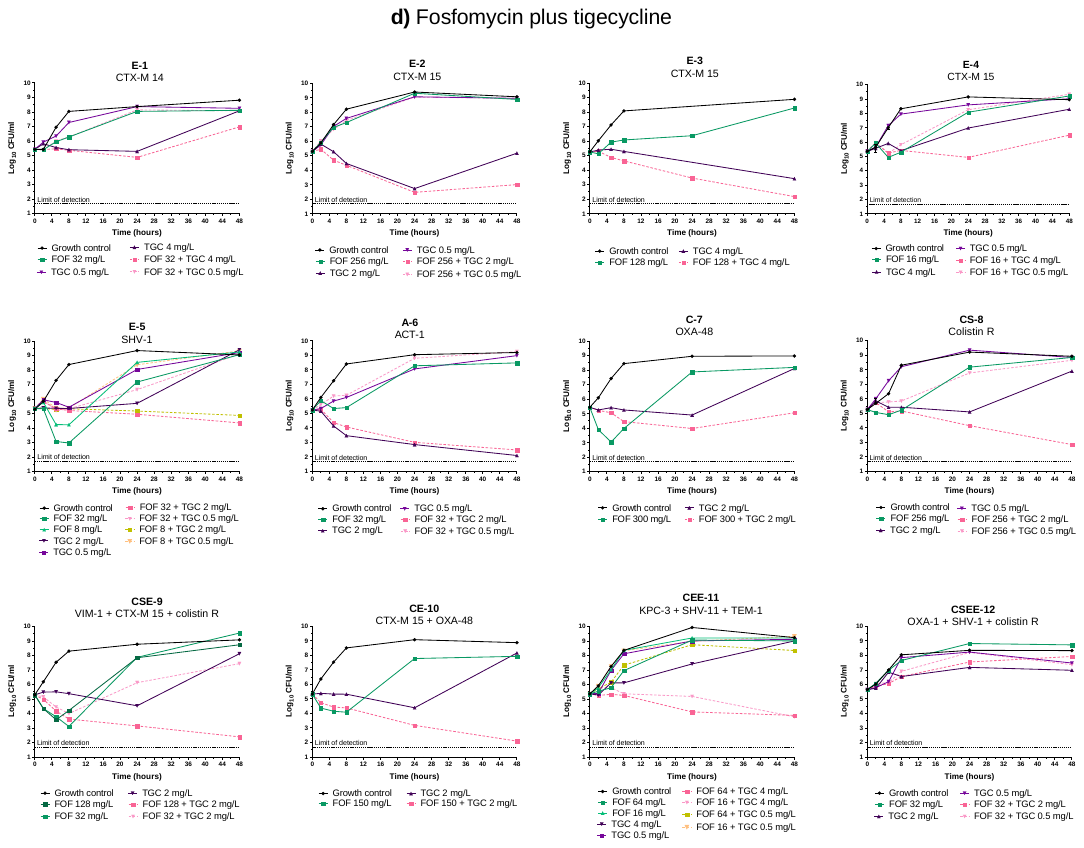

**Figure S2.** **Time-kill assays of azithromycin combined with last-line antibiotics against twelve *K. pneumoniae* isolates. (a)** Azithromycin plus fosfomycin; **(b)** Azithromycin plus colistin; **(c)** Azithromycin plus tigecycline.

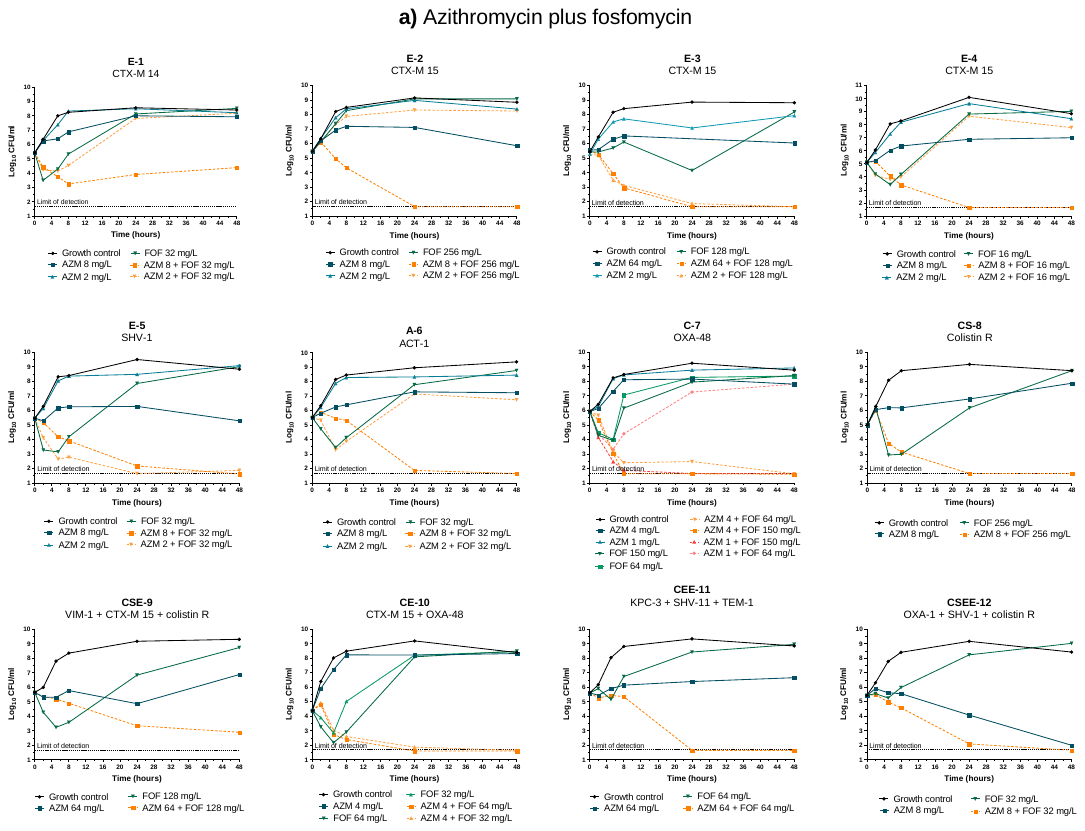

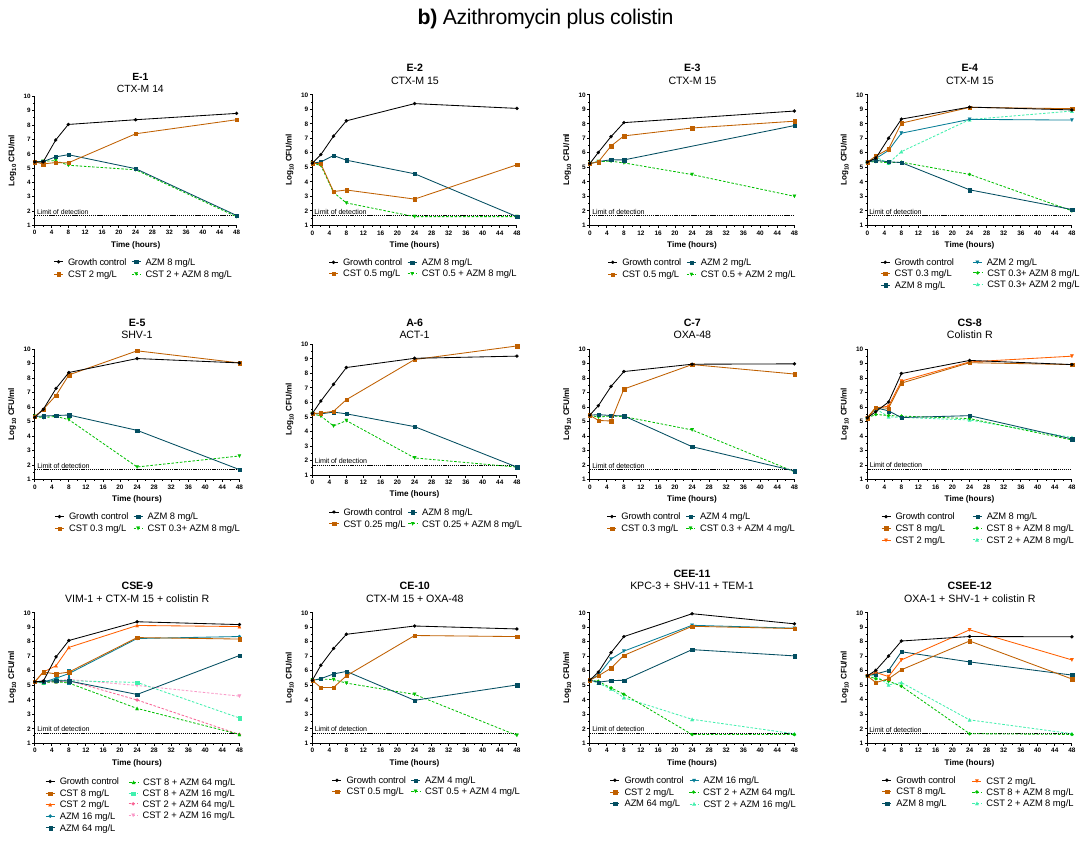

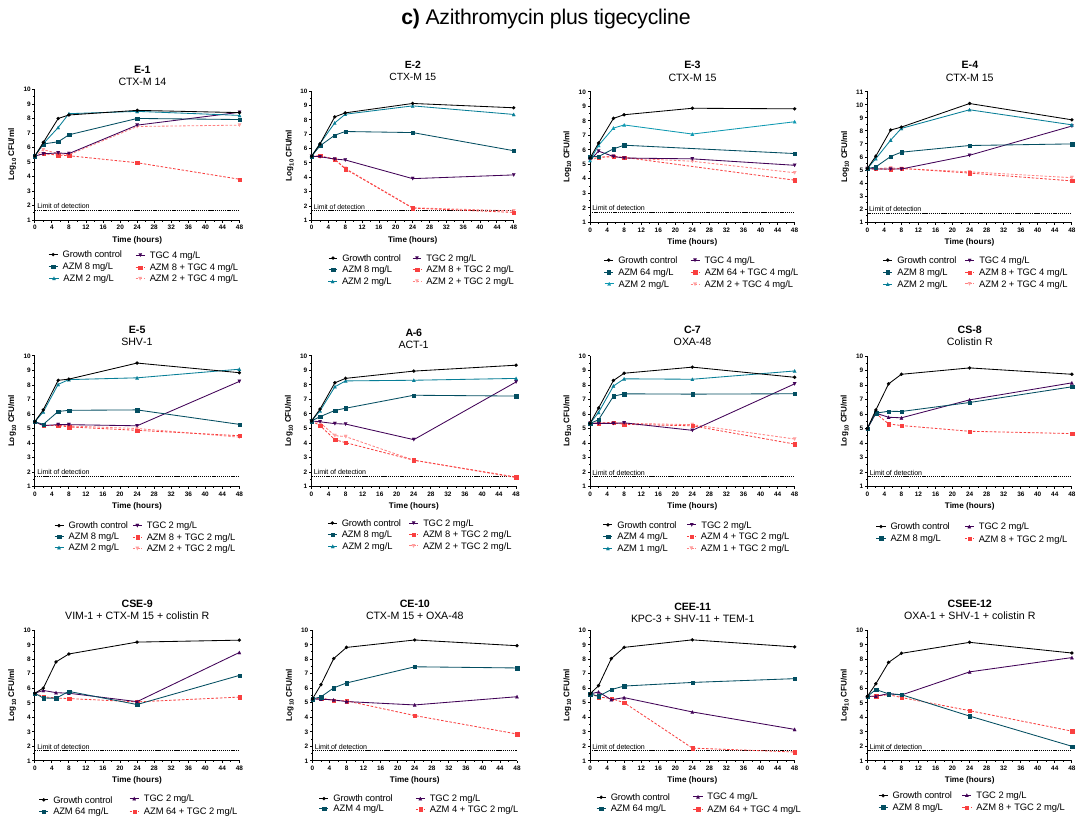
