## Supplementary Table S1 for "Repurposing azithromycin in combination with last-line fosfomycin, colistin and tigecycline against Multi-Drug Resistant *Klebsiella pneumoniae*"

**Table S1. Strain characterization and antimicrobial susceptibility for the twelve MDR/XDR K. pneumoniae isolates**

|  |  |  |  | **MIC (mg/L)^b^** | | | | | | | | | | | | | | | | | | | | |
| --- | --- | --- | --- | --- | --- | --- | --- | --- | --- | --- | --- | --- | --- | --- | --- | --- | --- | --- | --- | --- | --- | --- | --- | --- |
| **Isolate** | **Resistance mechanism** | **Specimen source** | **MDR/XDR classification^a^** | **AMK** | **GEN** | **TOB** | **AMP/AMX** | **AMC** | **TZP** | **FOX** | **CXM** | **CTX** | **CAZ** | **FEP** | **ATM** | **IPM** | **ETP** | **MEM** | **CIP** | **LVX** | **FOF** | **CST** | **TGC** | **SXT** |
| E-1 | CTX-M 14 | Rectal swab | XDR | ≤8 | >8 | ≤2 | >16 | >16/8 | >64 | >16 | >16 | >32 | >16 | >16 | >16 | 8 | >1 | 8 | >2 | 4 | ≤32 | ≤2 | >2 | >4/76 |
| E-2 | CTX-M 15 | Blood | MDR | ≤8 | ≤2 | >8 | >16 | >16/8 | >64 | >16 | >16 | >32 | >16 | >16 | >16 | ≤1 | >1 | 2 | >2 | >4 | >64 | ≤2 | ≤1 | ≤2/38 |
| E-3 | CTX-M 15 | Abscess | MDR | ≤8 | >8 | >8 | >16 | >16/8 | 64 | >16 | >16 | >32 | >16 | >16 | >16 | ≤1 | >1 | 2 | >2 | 2 | ≤32 | ≤2 | ≤1 | >4/76 |
| E-4 | CTX-M 15 | Blood | MDR | 16 | >4 | >4 | >16 | >32 | 16 | >16 | >8 | >32 | 32 | >8 | >4 | ≤1 | ≤0.12 | ≤0.12 | >1 | >1 | ≤16 | ≤2 | >2 | >4/76 |
| E-5 | SHV-1 + porin loss | Blood | MDR | ≤8 | ≤2 | ≤2 | >16 | >16/8 | >64 | 16 | ≤4 | ≤1 | ≤1 | 4 | ≤1 | ≤1 | ≤0.5 | ≤1 | ≤0.5 | ≤1 | ≤32 | ≤2 | ≤1 | ≤2/38 |
| A-6 | AmpC ACT-1 | SEIMC CCS07 | MDR | >32 | >8 | >8 | >16 | >16/8 | 64 | >16 | >16 | 32 | >16 | ≤1 | >16 | ≤1 | >1 | ≤1 | 2 | ≤1 | ≤32 | ≤2 | ≤1 | >4/76 |
| C-7 | OXA-48 | Blood | MDR | ≤8 | ≤2 | ≤2 | >16 | >16/8 | >64 | ≤8 | 8 | ≤1 | ≤1 | ≤1 | ≤1 | 4 | >1 | 4 | ≤0.5 | ≤1 | 64 | ≤2 | ≤1 | ≤2/38 |
| CS-8 | Colistin R | Urine | MDR | ≤8 | ≤2 | ≤2 | >16 | >16/8 | >64 | >16 | 16 | ≤1 | ≤1 | 4 | ≤1 | ≤1 | ≤0.5 | ≤1 | ≤0.5 | ≤1 | >64 | >4 | 2 | >4/76 |
| CE-9 | VIM-1 + CTX-M 15 + colistin R | SEIMC CCS04 | XDR | 16 | >8 | >8 | >16 | >16/8 | >64 | >16 | >16 | >32 | >16 | >16 | >16 | 2 | >1 | 8 | >2 | >4 | ≤32 | >4 | 2 | >4/76 |
| CE-10 | CTX-M 15 + OXA-48 | Blood | MDR | ≤8 | >4 | >4 | >8 | >32 | >16 | ≤8 | >8 | >32 | 32 | >8 | >4 | 8 | >1 | 1 | >1 | >1 | 32 | ≤2 | ≤1 | >4/76 |
| CEE-11 | KPC-3 + SHV-11 + TEM-1 | SEIMC CCS05 | XDR | 32 | 4 | >8 | >16 | >16/8 | >64 | >16 | >16 | >32 | >16 | >16 | >16 | >8 | >1 | >8 | >2 | >4 | 64 | >4 | >2 | >4/76 |
| CSEE-12 | OXA-1 + SHV-1 + colistin R | EARS QC | MDR | >32 | >8 | >8 | >16 | >16/8 | >64 | ≤8 | >16 | >32 | ≤1 | >16 | ≤1 | ≤1 | >1 | ≤1 | >2 | >4 | ≤32 | >4 | 2 | >4/76 |

^a^MDR/XDR categorization according to Magiorakos et al. (29): MDR: non-susceptible to ≥1 agent in ≥3 antimicrobial categories; XDR: non-susceptible to ≥1 agent in all but ≤2 categories

^b^MIC determined by automated broth microdilution (Microscan Walkaway®, Beckman Coulter, Spain) and clinical interpretation according to the corresponding EUCAST guidelines on the isolation date. Values in green were interpreted as "Susceptible", in dark yellow as "Susceptible, increased exposure" (EUCAST 2019), in light yellow as "Intermediate" and in red as "Resistant".

| AMK | Amikacin |
| --- | --- |
| GEN | Gentamicin |
| TOB | Tobramicin |
| AMP/AMX | Ampicillin/amoxicillin |
| AMC | Amoxicillin-clavulanate |
| TZP | Piperacillin-tazobactam |
| FOX | Cefoxitin |
| CXM | Cefuroxime |
| CTX | Cefotaxime |
| CAZ | Ceftazidime |
| FEP | Cefepime |
| ATM | Aztreonam |
| IPM | Imipenem |
| ETP | Ertapenem |
| MEM | Meropenem |
| CIP | Ciprofloxacin |
| LVX | Levofloxacin |
| FOF | Fosfomycin |
| CST | colistin |
| TGC | Tigecycline |
| SXT | Trimethoprim/sulfamethoxazole |
